# Pre-existing tissue mechanical hypertension at adherens junctions disrupts apoptotic extrusion in epithelia

**DOI:** 10.1101/2023.08.31.555652

**Authors:** Zoya Mann, Fayth Lim, Suzie Verma, Bageshri N. Nanavati, Julie M. Davies, Jakob Begun, Edna C. Hardeman, Peter W. Gunning, Deepa Subramanyam, Alpha S. Yap, Kinga Duszyc

## Abstract

Apical extrusion is a tissue-intrinsic process that allows epithelia to eliminate unfit or surplus cells. This is exemplified by the early extrusion of apoptotic cells, which is critical to maintain the epithelial barrier and prevent inflammation. Apoptotic extrusion is an active mechanical process, which involves mechanotransduction between apoptotic cells and their neighbours, as well as local changes in tissue mechanics. Here we report that the pre-existing mechanical tension at adherens junctions conditions the efficacy of apoptotic extrusion. Specifically, increasing baseline mechanical tension by overexpression of a phosphomimetic Myosin II regulatory light chain (MRLC) compromises apoptotic extrusion. This occurs when tension is increased in either the apoptotic cell or its surrounding epithelium. Further, we find that the pro-inflammatory cytokine, TNFα, stimulates Myosin II and increases baseline AJ tension to disrupt apical extrusion, causing apoptotic cells to be retained in monolayers. Importantly, reversal of mechanical tension with an inhibitory MRLC mutant or tropomyosin inhibitors is sufficient to restore apoptotic extrusion in TNFα-treated monolayers. Together, these findings demonstrate that baseline levels of tissue tension are important determinants of apoptotic extrusion, which can potentially be co-opted by pathogenetic factors to disrupt the homeostatic response of epithelia to apoptosis.

## Introduction

Epithelia constitute primary body barriers that protect their underlying tissues from environmental dangers. Epithelial cells are physically coupled to one another, forming tissue layers that can respond to diverse stimuli in a well-coordinated manner (Lecuit and Yap 2015). Adherens junctions (AJs) are specialized epithelial cell-cell junctions, built upon classical cadherin adhesion complexes that physically couple to the contractile actomyosin network. This cytoskeletal association can generate mechanical tension in the junctions, creating the potential for AJs to form a tensile network that connects cells together in epithelial monolayers (Charras and Yap 2018). Dynamic patterns of change in AJ tension influence cell-cell rearrangements during morphogenesis in the embryo (Heer and Martin 2017). They also participate in homeostasis during post-developmental life, as exemplified by the epithelial response to apoptosis (Lubkov and Bar-Sagi 2014; Michael et al. 2016; Duszyc, Gomez, et al. 2021).

Epithelial integrity is constantly challenged by environmental stresses that can cause apoptosis. Extrusion constitutes a first-line response to the apoptotic challenge that robustly eliminates dying cells in a manner of minutes (Duszyc, Gomez, et al. 2021; Rosenblatt, Raff, and Cramer 2001; Michael et al. 2016). If the extrusion response fails, retained apoptotic cells can disrupt the functional epithelial barrier and provoke pro-inflammatory responses from neighbouring epithelial cells (Duszyc et al. 2023). Successful extrusion involves a complex series of mechanical changes in the apoptotic cells and their proximal neighbours (Fig S1A) (Teo, Tomatis, et al. 2020). In one set of changes, hypercontractility of apoptotic cells stimulates the adjacent cells to assemble a contractile cortex at their interface with the apoptotic cells (Andrade and Rosenblatt 2011; Atieh, Ruiz, and Eisenhoffer 2021; Duszyc, Gomez, et al. 2021; Lubkov and Bar-Sagi 2014). Contraction of this cortical network drives the physical expulsion of the dying cells. In parallel, AJ in neighbour cells that are oriented tangential to the apoptotic cell undergo mechanical relaxation (Teo, Tomatis, et al. 2020). This facilitates apoptotic extrusion, and itself appears to be a response to transient early relaxation in the acutely injured cells (Teo, Tomatis, et al. 2020). These complex local patterns of mechanical change therefore appear to reflect multiple mechanotransduction pathways that mediate communication from apoptotic cells to their neighbours.

The dynamic changes in AJ tension associated with apoptotic extrusion occur upon a pre-existing or baseline pattern of mechanical junctional tension (Teo, Tomatis, et al. 2020). This raises the interesting question of whether aberrant change in baseline AJ tension can affect the efficacy of apoptotic extrusion. Recently, we reported that an increase in baseline AJ tension (mechanical hypertension, for short) compromised the ability of epithelial monolayers to eliminate oncogene-expressing cells by apical extrusion (Teo, Gomez, et al. 2020). Whether this may apply for apoptotic extrusion is not yet known. We now report that increasing pre-existing AJ tension using a phosphomimetic Myosin regulatory light chain (MRLC) mutant (MRLC-DD) decreases the ability of epithelial monolayers to execute apoptotic extrusion. Further, we show that the inflammatory cytokine, TNFα, also increases baseline AJ tension, and this compromises the elimination of apoptotic cells by apical extrusion. This highlights how baseline tissue tension regulates apical extrusion and suggests that changes in pre-existing tension may cause homeostasis to be compromised in disease.

## Results and Discussion

### MRLC-DD increases adherens junction tension in Caco-2 monolayers

To assess the potential implications of altered epithelial mechanics, we established a Caco-2 line stably expressing a phosphomimetic mutant of Myosin Regulatory Light Chain (MRLC-DD) (Fig S1B,C). As actomyosin is the major contributor to cellular force generation, MRLC-DD is predicted to enhance actomyosin activity, resulting in increased intercellular junctional tension.

To characterize the impact of MRLC-DD expression on AJ tension, we first examined the cellular localisation of non-muscle Myosin IIA and IIB, as well as the F-actin network. Levels of NMIIA and Myosin IIB (Fig 1A-C) and F-actin (Fig 1D-E) were increased at AJ in the MRLC-DD-expressing monolayer when compared to control GFP-expressing cells. The increase in junctional Myosin IIA and IIB was not associated with overall changes in expression levels of these proteins (Fig S1D-G). This suggested that contractile tension might be increased at AJ in MRLC-DD-expressing cells.

**Figure 1.**
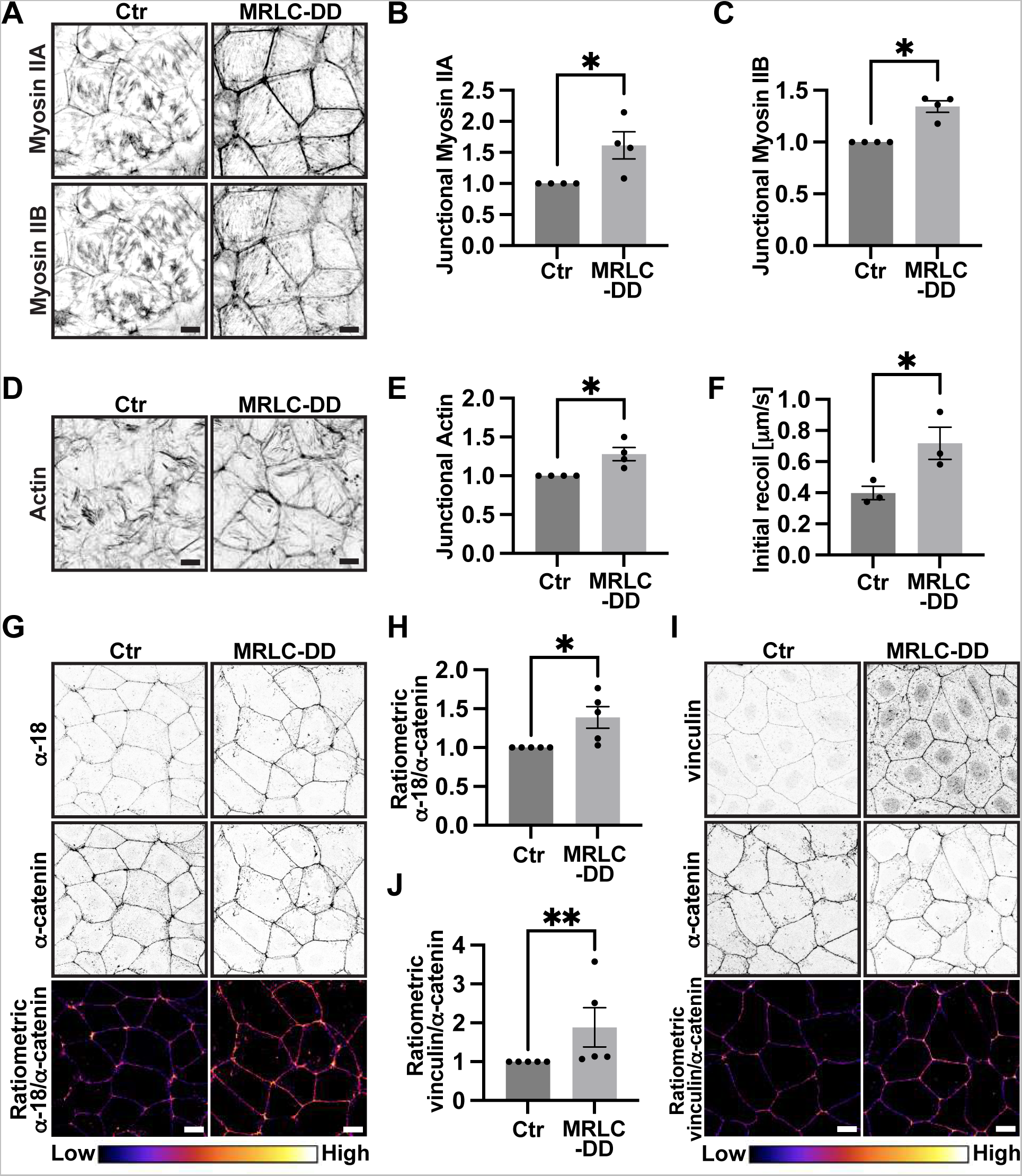
MRLC-DD increases junctional tension in Caco-2 monolayers. **(A-C)** Representative images (A) and quantification of Myosin IIA (B) and IIB (C) junctional localisation in control and MRLC-DD cell lines. **(D-E)** Representative images (D) and quantification (E) of actin junctional localisation in control and MRLC-DD cell lines. **(F)** Initial recoil of ablated junctions in control and MRLC-DD cell lines. **(G-H)** Representative images (G) and quantification (H) of junctional localisation of α-catenin in open conformation (α-18) and total α-catenin in control and MRLC-DD cell lines. Ratiometric images represent junctional intensity of α-18 divided by junctional intensity of total α-catenin. **(I-J)** Representative images (I) and quantification (J) of junctional localisation of vinculin and α-catenin in control and MRLC-DD cell lines. Ratiometric images represent junctional intensity of vinculin divided by junctional intensity of total α-catenin. Scale bars: 15μm. XY panels are maximum projection views of all z-stacks. All data are means ± SEM; *p< 0.05, **p< 0.01, calculated from n≥3 independent experiments analysed with unpaired Student’s t-test.

To pursue this further, we applied more direct approaches to evaluate potential changes in AJ tension. First, we tested whether MRLC-DD affects junctional tension by assessing the recoil of junctions (visualised by mCherry tagged endogenous ZO-1) after they were cut with a 2-photon laser beam (Liang, Michael, and Gomez 2016). Following laser ablation, junctions in the GFP-expressing control cells exhibited a prompt recoil (Fig S1H-J), indicating that junctions of steady-state Caco-2 monolayers were tensile. Importantly, the recoil (Fig S1H-J) and the initial velocity of the recoil (Fig 1F) was significantly increased in the MRLC-DD-expressing cells, consistent with increased junctional tension. To complement this experiment, we asked whether we could detect tension-associated changes at a molecular level. Specifically, we focused on cadherin-associated α-catenin, which is reported to undergo conformational changes upon application of tension to the E-cadherin complex (Noordstra, Morris, and Yap 2023; Yonemura et al. 2010). Here, we utilised an antibody (α-18 mAb), which detects a tension-exposed epitope in the central, M-domain of α-catenin (Nagafuchi and Tsukita 1994). To correct for heterogeneity in staining intensity among samples, we used the ratio of α-18 to total α-catenin as a readout of molecular-level tension. MRLC-DD-expressing cells displayed a significantly higher ratio of junctional α-18/total α-catenin than the control cells (Fig 1G-H). We corroborated this by measuring junctional levels of vinculin, which is recruited to α-catenin under tension (Yonemura et al. 2010; Yao et al. 2014). Consistent with our earlier results, junctional vinculin, corrected for α-catenin levels, was increased by expression of MRLC-DD (Fig 1I-J). Taken together, our observations confirmed that junctional tension was elevated by expression of the MRLC-DD transgene in Caco-2 monolayers, which is consistent with previous reports.

### Elevated baseline adherens junctional tension disrupts apoptotic extrusion

Then, we asked whether global increase in baseline junctional tension could affect the ability of epithelia to expel apoptotic cells. To test this idea, we treated control and MRLC-DD-expressing monolayers with the topoisomerase II inhibitor etoposide, which is commonly used to induce sporadic apoptosis in epithelial cell culture models (Duszyc, Gomez, et al. 2021; Michael et al. 2016; Teo, Tomatis, et al. 2020; Andrade and Rosenblatt 2011). Strikingly, while control cells were able to robustly expel cleaved caspase-3 positive, apoptotic cells (Fig 2A-B), the MRLC-DD cells were significantly less efficient at eliciting extrusion (Fig 2A-B). We then induced apoptosis by microirradiating nuclei with a 2-photon laser (Duszyc, Gomez, et al. 2021; Teo, Tomatis, et al. 2020; Duszyc, Noordstra, et al. 2021); this provided tight temporal control of injury to monitor the extrusion process with high spatio-temporal resolution (Fig 2C). To visualize the behaviour of the neighbour cells we performed these experiments in control and MRLC-DD-expressing cell monolayers in which endogenous ZO-1 was tagged with mCherry by CRISPR/Cas9 genome editing (Duszyc et al. 2023). We observed that AnnexinV-positive apoptotic cells were successfully extruded apically out of the control monolayer, typically within less than 60min following laser injury (Fig 2C-E). This was associated with elongation of the immediate neighbours of the apoptotic cells, culminating in formation of a rosette-like pattern underneath the apically extruded cells (Fig 2C). In contrast, MRLC-DD-expressing monolayers commonly failed to extrude the dying cells (Fig 2C-E). Instead, AnnexinV-positive apoptotic cells persisted within the epithelial layers for over 60min (Fig 2E). Together, these observations indicated that a global increase of baseline AJ tension can impede the epithelial-intrinsic ability of monolayers to extrude apoptotic cells.

**Figure 2.**
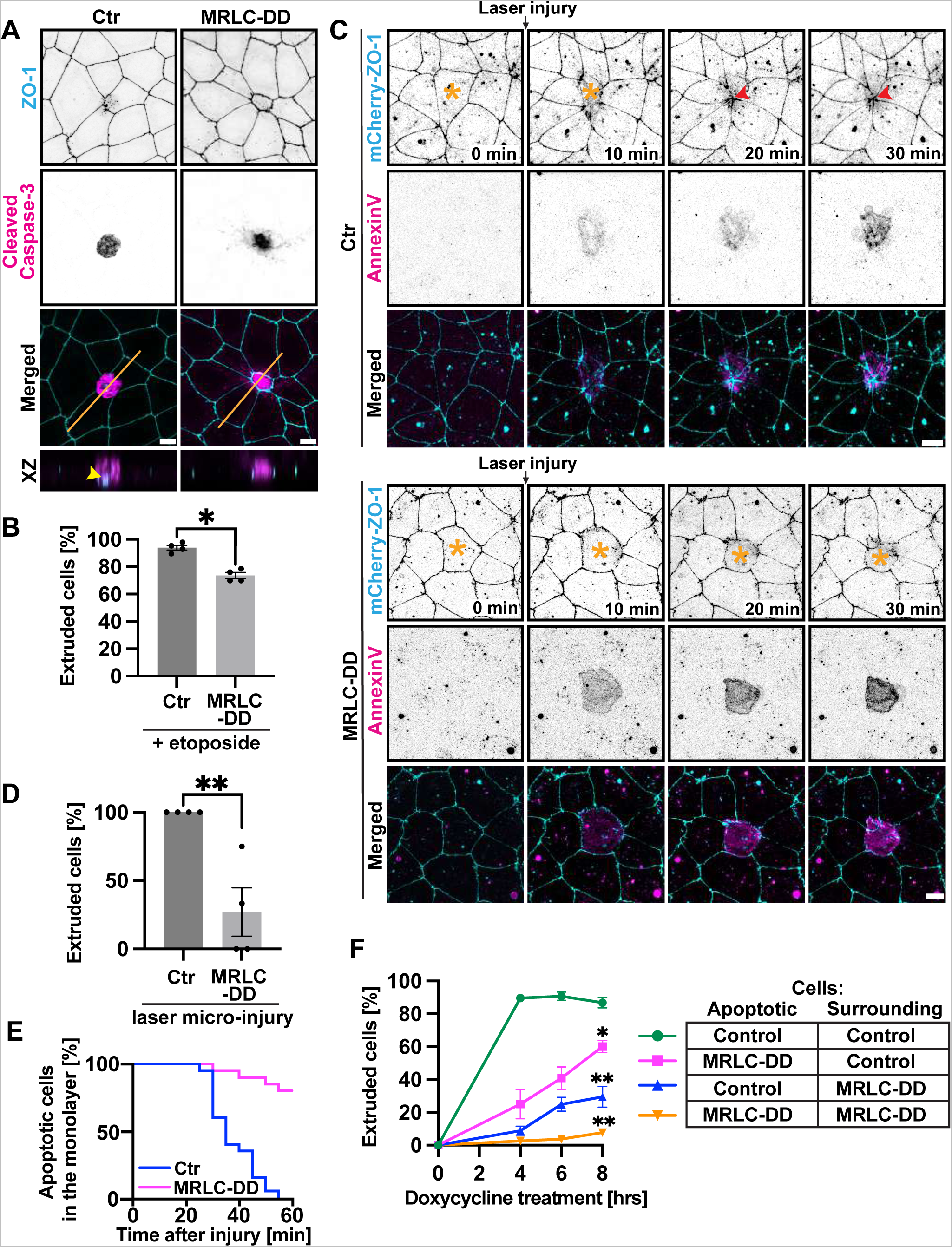
Increased baseline junctional tension compromises apoptotic extrusion. **(A-B)** Representative images (A) and quantification (B) of extrusion efficiency in etoposide treated (500μM, 8hrs) control and MRLC-DD cell lines. Orange lines-location of the XZ views; arrowhead - junctional closure underneath extruded apoptotic cell. **(C-E)** Selected frames from live imaging of 2-photon laser-mediated apoptosis in control and MRLC-DD cell lines (C). Apoptosis was confirmed by positive AnnexinV staining. Asterisks– apoptotic cells in the monolayers; arrowheads – junctional closure underneath extruded apoptotic cell. (D) Quantification of efficiency of apoptotic extrusion in control and MRLC-DD cell lines. Extrusion was classified as successful if apoptotic cells were expelled from the monolayers within 60min after the 2-photon laser micro-injury. (E) Representation of time [min] apoptotic cells were present within the monolayer before extrusion was completed. (F) Quantification of extrusion efficiency of PUMA-expressing apoptotic cells at 4, 6 and 8hrs of doxycycline (2μg/ml) treatment in mixed cell cultures. Scale bars: 15μm. XY panels are maximum projection views of all z-stacks. All data are means ± SEM; *p< 0.05, **p< 0.01, calculated from n≥3 independent experiments analysed with unpaired Student’s t-test (B, D) or two-way ANOVA (F).

These results complement our earlier report that enhanced tissue tension compromises the efficiency of oncogenic extrusion (Teo, Gomez, et al. 2020). Of note, oncogenic extrusion was inhibited when tension was upregulated exclusively in the surrounding epithelium, rather than in the oncogene-expressing cell itself. We then asked whether this holds true for apoptotic extrusion. For this, we employed a cell-mixing protocol that allowed us to independently increase contractile tension in these different cell populations (Fig S2A). To induce apoptosis, we used the Tet-on system to express PUMA, a BH3-only protein that has been reported to robustly elicit cell death in epithelial cells (Duszyc, Gomez, et al. 2021; Bonfim-Melo et al. 2022). First, we generated a stable cell line expressing PUMA^Tet- on^ that was later transduced with a lentivirus carrying either MRLC-DD or a control soluble GFP transgene (MRLC-DD^PUMA^ and Control^PUMA^, respectively) (Fig S2A) (Duszyc, Gomez, et al. 2021). Then, we mixed the MRLC-DD^PUMA^ or Control^PUMA^ cells with control or MRLC-DD cells in a 1:100 ratio, so that small clusters of PUMA-expressing cells (typically <2-4) were surrounded by non-apoptotic cells (Fig S2A). The mixed cultures were grown to confluency and expression of PUMA was induced with doxycycline. As expected, Control^PUMA^ cells were efficiently extruded by surrounding control neighbours (Fig 2F, Fig S2B), while the hypertensile MRLC-DD^PUMA^ cells were retained within the equally tensile MRLC-DD neighbouring epithelium (Fig 2F, Fig S2B).

In contrast, the rates of extrusion were significantly decreased when pre-existing tension was increased in either the surrounding epithelium or in the apoptotic cells themselves (Fig 2F, Fig S2B). Increasing mechanical tension in these different cell populations therefore appeared to independently compromise apoptotic extrusion. The reason for this remains to be elucidated. One possibility for the surrounding epithelium is that mechanical hypertension prevents neighbouring cells from relaxing their orthogonal junctions (Fig S1A) sufficiently to elongate and mediate extrusion (Teo, Tomatis, et al. 2020). From the perspective of the cells that are targeted for apoptosis, early apoptotic cells were reported to release tension before they become hypercontractile. This initial relaxation of apoptotic cells appeared to generate a signal for neighbour cells to elongate their orthogonal junctions (Teo, Tomatis, et al.). Increasing pre-existing contractility in apoptotic cells may then prevent them generating a sufficient signal to engage elongation by their neighbours. These are interesting questions for future research.

### TNFα upregulates junctional tension

Can AJ hypertension and retention of apoptotic cells occur in a patho-physiological context? To address this question, we focused on TNFα, a key pro-inflammatory cytokine upregulated in many inflammatory and autoimmune diseases, even when they are in clinical remission (Ruder, Atreya, and Becker 2019; Jang et al. 2021; Bradley 2008). Importantly, TNFα has been previously reported to transcriptionally upregulate Myosin Light Chain Kinase (MLCK), which can directly activate non-muscle Myosin II (Graham et al. 2006; Ma et al. 2005). Therefore, we tested whether treatment of Caco-2 cells with TNFα affects AJ tension.

We found that TNFα-treated cells displayed more junctional Myosin IIA than untreated control cells (Fig 3A-B). Also, the junctional pool of phosphorylated MRLC (pMRLC) was significantly increased by TNFα (Fig 3C-D). This was accompanied by an increase in baseline AJ tension, as measured by ratiometric α-18 mAb/total α-catenin imaging (Fig 3E-F). Additionally, both the recoil (Fig 3G, Fig S3A-B) and the initial speed of junctional recoil (Fig 3H) were enhanced in TNFα-treated monolayers. Together, the results indicate that stimulation of NMII by TNFα increases the mechanical tension in AJ.

**Figure 3.**
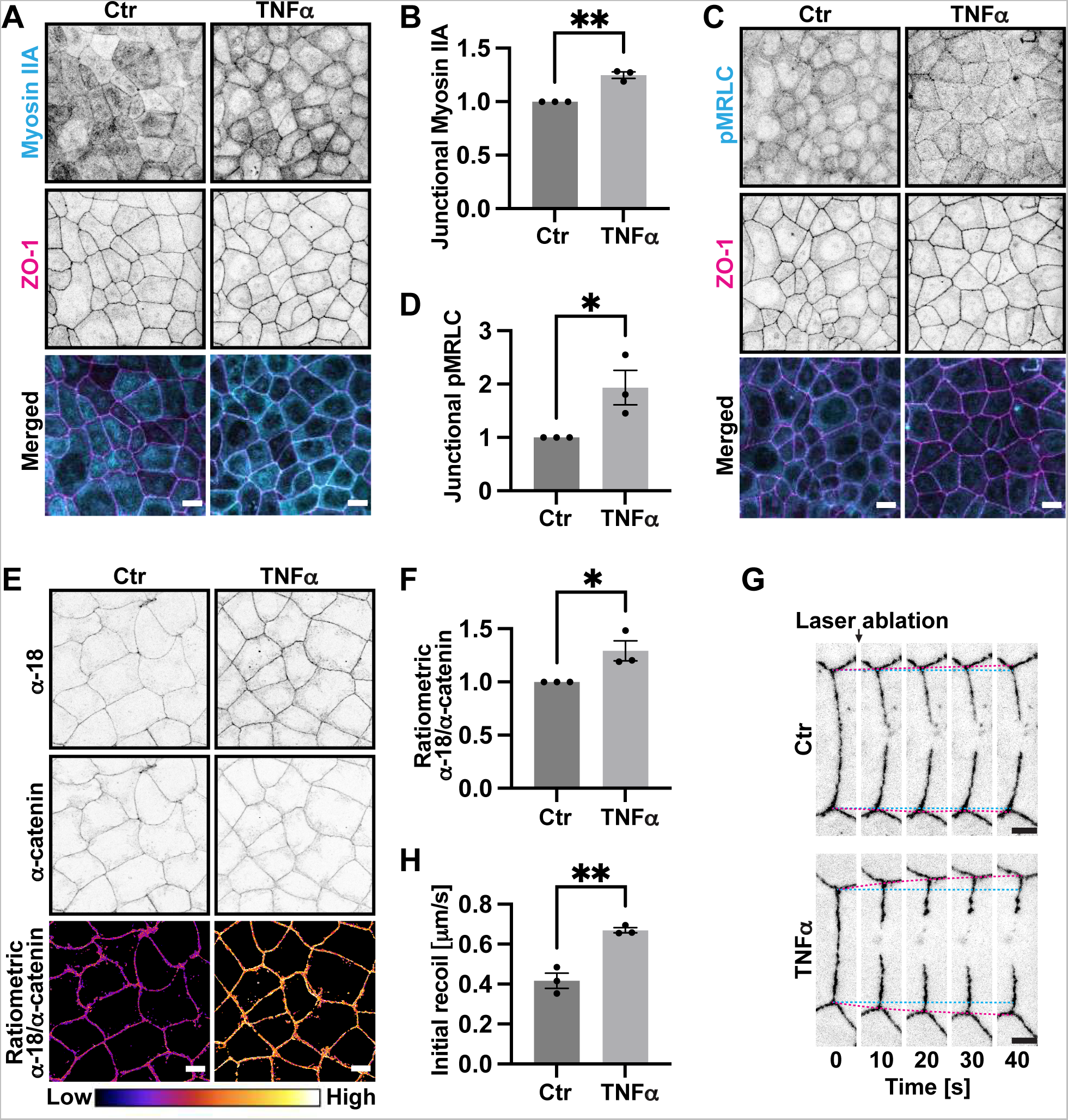
TNFα upregulates junctional tension. **(A-B)** Representative images (A) and quantification (B) of Myosin IIA junctional localisation in control and TNFα-treated cell lines. **(C-D)** Representative images (C) and quantification (D) of phosphorylated MRLC (pMRLC) junctional localisation in control and TNFα-treated cell lines. **(E-F)** Representative images (E) and quantification (F) of junctional localisation of α-catenin in open conformation (α-18) and total α-catenin in control and TNFα-treated cell lines. Ratiometric images represent junctional intensity of α-18 divided by junctional intensity of total α-catenin. **(G-H)** Selected frames (G) from live imaging of 2-photon laser-mediated junction ablation followed by junctional recoil in control and TNFα-treated cells. Calculated speed of initial recoil (H) of ablated junctions in control and TNFα-treated cells. Scale bars: 15μm. XY panels are maximum projection views of all z-stacks. All data are means ± SEM; *p<0.05, **p<0.01, calculated from n≥3 independent experiments analysed with unpaired Student’s t-test.

### TNFα perturbs apoptotic extrusion

Then, we asked if induction of mechanical hypertension by TNFα affected the ability of epithelia to eliminate apoptotic cells by apical extrusion. To test this, we induced etoposide-mediated apoptosis in control and TNFα-treated cellular monolayers and quantified the percentage of apoptotic cells that were successfully extruded (Fig 4A, Fig S4A). Whilst control cells robustly expelled almost 100% of apoptotic AnnexinV-positive cells, TNFα treatment decreased the extrusion efficiency to less than 50% (Fig 4A, Fig S4A). We considered the possibility that the increased number of retained apoptotic cells we saw was due to enhanced cell mortality, as TNFα TNFR1 receptor is capable of inducing apoptosis (Micheau and Tschopp 2003). To address this, we live-imaged TNFα-treated and control cell cultures. However, we did not detect a significant difference in the number of cells that underwent apoptosis in these two groups (Fig S4B). As expected, etoposide increased the number of apoptotic cells detected during live-imaging, but the incidence of apoptosis was comparable between control and TNFα-treated monolayers (Fig S4B). Consistent with this, we did not detect cleaved caspase-3 immunoblot bands in control or TNFα-treated cells (Fig S4C-D), and the intensity of the bands was comparable in these two groups upon etoposide treatment (Fig S4C-D). Therefore, we reasoned that TNFα treatment did not stimulate apoptosis, nor did it sensitise cells to etoposide-induced cell death, in our system.

**Figure 4.**
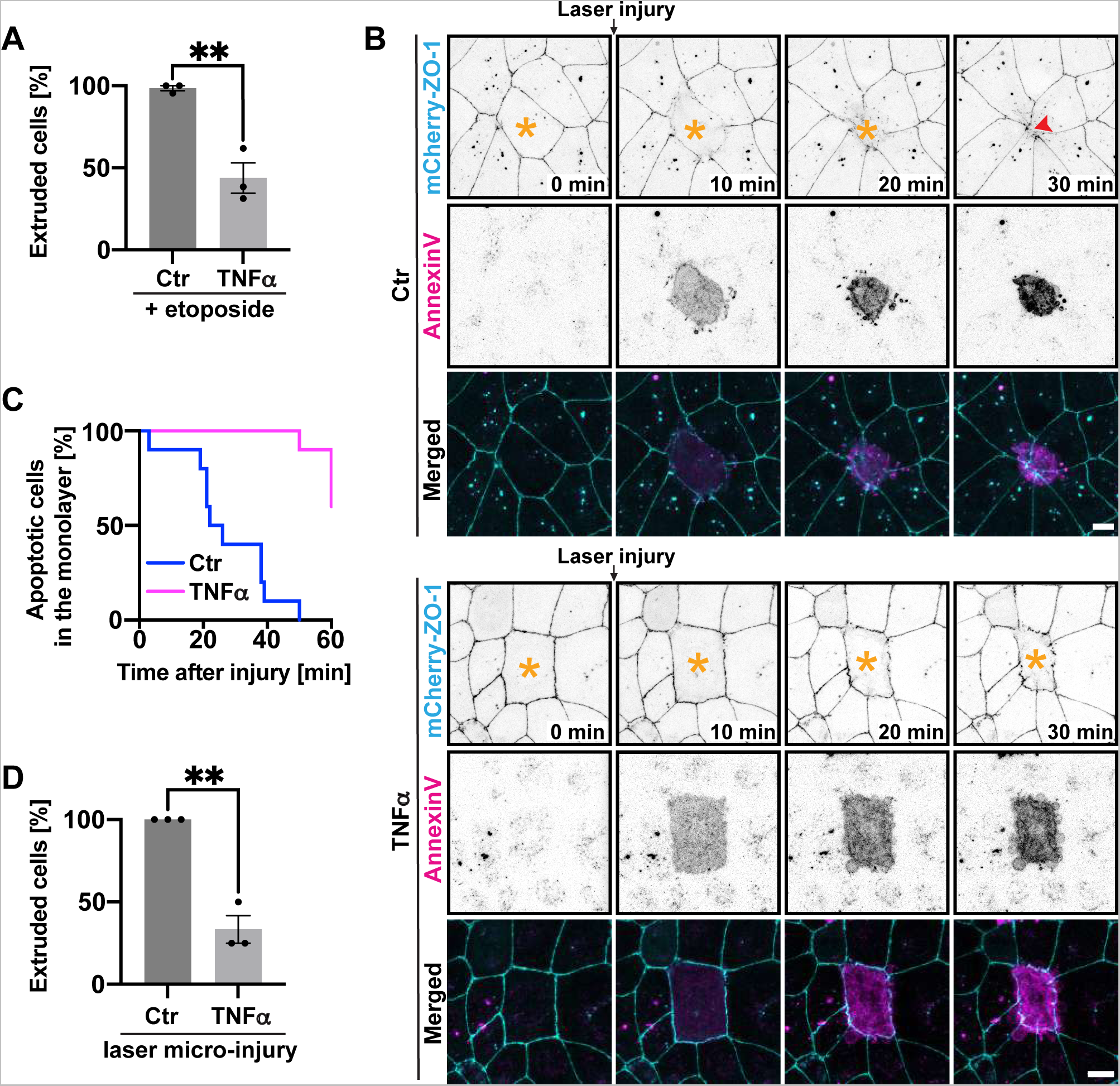
TNFα impairs apoptotic extrusion. **(A)** Quantification of extrusion efficiency in control and TNFα-treated cell lines. Apoptosis was induced by etoposide treatment (500μM, 8hrs). **(B-D)** (B) Selected frames from live imaging of 2-photon laser-mediated apoptosis in control and TNFα-treated cell lines. Apoptosis was confirmed by positive AnnexinV staining. Asterisks– apoptotic cells in the monolayer; arrowhead – junctional closure underneath extruded apoptotic cell. (C) Representation of time [min] apoptotic cells are present within the monolayer before extrusion is completed. (D) Quantification of efficiency of apoptotic extrusion in control and TNFα-treated cell lines; extrusion was classified as successful if apoptotic cells were expelled from the monolayers within 60min after the 2-photon laser micro-injury. Scale bars: 15μm. XY panels are maximum projection views of all z-stacks. All data are means ± SEM; **p<0.01 calculated from n≥3 independent experiments analysed with unpaired Student’s t-test (A, D).

We inferred instead that TNFα treatment perturbed extrusion of apoptotic cells. We tested this directly by inducing apoptosis with 2-photon nuclear microirradiation. In line with our previous observations, control monolayers expelled apoptotic cells typically within 30 min post injury (Fig 4B-C). In contrast, TNFα-treated cells were frequently unable to extrude apoptotic cells even 60 min post injury, leading to prolonged retention of cell corpses within the epithelial layers (Fig 4B-D). Together, this reinforced the notion that TNFα perturbs the epithelial-intrinsic ability to extrude apoptotic cells.

### Correction of exacerbated tension rescues apoptotic extrusion in TNFα-treated monolayers

As TNFα is known to cause diverse cellular effects (Tracey and Cerami 1994; Gough and Myles 2020; Leppkes et al. 2014), we then asked whether the disruption of apoptotic extrusion in TNFα-treated monolayers was indeed caused by mechanical hypertension. For this we expressed a phosphodeficient mutant of MRLC (MRLC-AA) to reduce AJ tension in TNFα-treated monolayers. We confirmed that MLRC-AA reversed enhanced baseline junctional tension, as measured by ratiometric α-18 mAb/total α-catenin immunostaining (Fig 5A-B). Importantly, we found that MRLC-AA significantly increased the percentage of apoptotic cells that were apically expelled from TNFα-treated epithelial monolayers (Fig 5C-D). This implied that correcting junctional hypertension could restore apoptotic extrusion to TNFα-treated monolayers.

**Figure 5.**
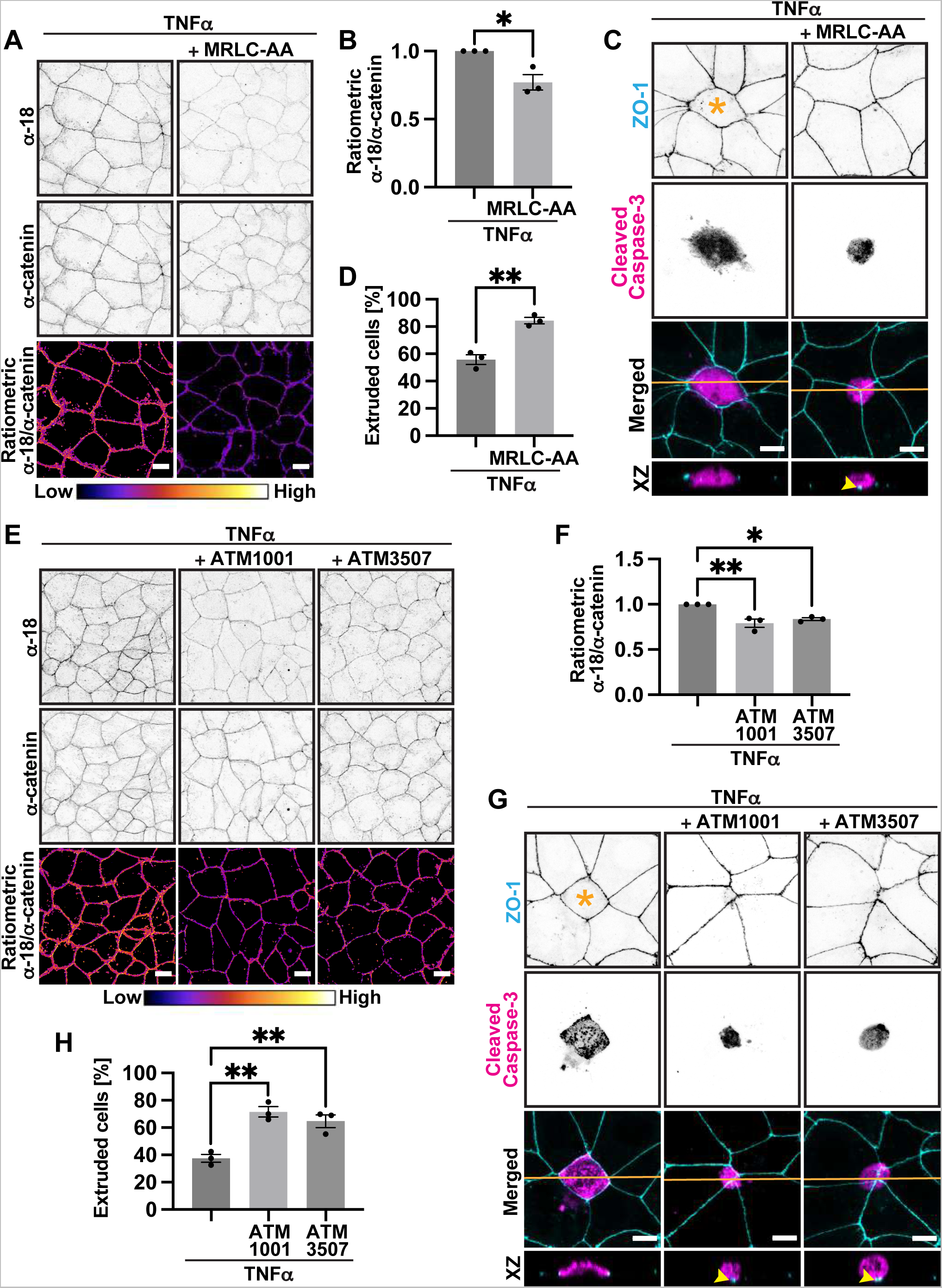
Correction of mechanical hypertension rescues apoptotic extrusion in TNFα–treated monolayers. **(A-B)** Representative images (A) and quantification (B) of junctional localisation of α-catenin in open conformation (α-18) and total α-catenin in TNFα-treated cell lines upon expression of MRLC-AA. Ratiometric images represent junctional intensity of α-18 divided by junctional intensity of total α-catenin. **(C-D)** Representative images (C) and quantification (D) of extrusion efficiency in TNFα-treated cells upon expression of MRLC-AA. Apoptosis was inducted by etoposide treatment (500μM, 8hrs). Orange lines - location of the XZ views; asterisk – retained apoptotic cell; arrowhead – junctional closure underneath the extruded apoptotic cells. **(E-F)** Representative images (E) and quantification (F) of junctional localisation of α-catenin in open conformation (α-18) and total α-catenin in TNFα stimulated cell lines upon treatment with tropomyosin inhibitors ATM1001 (2.5μM, 8hrs) and ATM3507 (2.5μM, 8hrs). Ratiometric images represent junctional intensity of α-18 divided by junctional intensity of total α-catenin. **(G-H)** Representative images (G) and quantification (H) of extrusion efficiency in TNFα-stimulated cells upon treatment with tropomyosin inhibitors ATM1001 (2.5μM, 8hrs) and ATM3507 (2.5μM, 8hrs). Apoptosis was inducted by etoposide treatment (500μM, 8hrs). Orange lines - location of the XZ views; asterisk – retained apoptotic cell; arrowheads – junctional closure underneath the extruded apoptotic cells. Scale bars: 15μm. XY panels are maximum projection views of all z-stacks. All data are means ± SEM; *p<0.05, **p<0.01, calculated from n≥3 independent experiments analysed with unpaired Student’s t-test (B, D) or one-way ANOVA (F, H).

As an independent test of this idea, we used the structurally distinct tropomyosin inhibitors (TPMi) ATM1001 and ATM3507 (Kee et al. 2018; Currier et al. 2017) to reverse AJ tension. Tropomyosin supports actomyosin to generate contractile tension at AJs and drug inhibition of tropomyosin or depletion of its Tm5NM1 (Tpm3.1) or Tm5NM2 (Tpm3.2) isoforms was previously reported to decrease junctional epithelial tension (Caldwell et al. 2014). We found that treatment with either ATM1001 or ATM3507 reversed AJ hypertension (Fig 5E-F) and increased the percentage of apoptotic cells that were extruded from the TNFα-treated monolayers (Fig 5G-H). Therefore, reversal of mechanical hypertension can restore the ability of Caco-2 monolayers to expel apoptotic cells despite the continuous presence of TNFα. This further indicated that TNFα perturbed apoptotic extrusion through induction of AJ hypertension.

## Concluding remarks

Together, these findings indicate that the pre-existing tension in epithelial monolayers can condition the efficacy of apical extrusion. Specifically, apoptotic extrusion was inhibited by baseline mechanical hypertension, when hypertension was induced by manipulating Myosin II or stimulation with TNFα. This carries the implication that pathogenetic factors, such as inflammatory cytokines, may exert some of their effects by altering tissue mechanics. For example, although TNFα induces apoptosis in some cells, this did not appear to be the case in our studies. The observed increase in number of apoptotic cells within monolayers was due to a failure of extrusion. Whether this might apply in other contexts is an interesting question. Furthermore, while apoptosis is widely regarded as an immunologically silent phenomenon, we have recently reported that epithelia activate an acute, IL-8 driven, inflammatory response when apical extrusion fails to eliminate apoptotic cells (Duszyc et al. 2023). This raises the interesting question of whether mechanical hypertension may increase the risk of provoking overt inflammation by blocking extrusion and causing apoptotic cells to be retained in epithelia.

## Materials and Methods

### Cell lines and culture

Caco-2 and HEK-293T cells were cultured in RPMI and DMEM media respectively, supplemented with 10% v/v Fetal Bovine Serum (FBS), 1% non-essential amino acids (NEAA), 1% L-glutamine, 100 U/ml Penicillin/Streptomycin and grown at 37°C in 5% CO2 atmosphere. To produce lentivirus particles, required pLL5.0 expression vectors and packaging vectors were transfected into HEK-293T cells using Lipofectamine 2000 (Invitrogen). 24 hrs after transfection cell medium was changed to fresh DMEM supplemented with 10% serum, 1% NEAA and 1% L-glutamine. 48hrs later virus containing medium was collected, fileted though a 0.45μm syringe-driven filters and subsequently concentrated on Amicon® Ultra-15 Centrifugal Filter Unit (Millipore, cat#UFC910096). Concentrated media containing lentiviral particles was used for infection of Caco-2 cells.

### Antibodies

Primary antibodies used in this study were as follows-

Rabbit pAb against Vinculin (Abcam, Cat#ab91459; RRID: AB_2050446)

Mouse mAb against α-catenin (Invitrogen, Cat# α-CAT-7A4)

Rat pAb against α-18 (a kind gift from Akira Nagafuchi, Nara Medical University, Japan)

Rabbit mAb against Cleaved caspase-3 (Cell Signalling Technologies, Cat#9664; RRID: AB_2070042)

Mouse mAb against ZO-1 (Invitrogen, Cat#33-9100)

Mouse mAb against NMIIA (Abcam, Cat#ab55456; RRID: AB_944320)

Rabbit mAb against NMIIB (Biolegend, Cat#909901)

Rabbit mAb against GAPDH (Abcam, Cat#ab181603; RRID: AB_2687666)

Mouse mAb against β-Tubulin (Cat#T4026; RRID: AB_477577)

Mouse mAb against GFP (Sigma, Cat#11814460001 Roche; RRID: AB_390913)

Rat mAb against E-cadherin ectodomain (Invitrogen, Cat#13-1900)

Alexa Fluor™ 647 Phalloidin (Invitrogen, Cat#A22287)

Secondary antibodies were species-specific conjugated with AlexaFluor-488, 546, 594 or 647 (Invitrogen) for immunofluorescence, or with horseradish peroxidase (Bio-Rad Laboratories) for immunoblowng.

### TNFα and ATM1001/ATM3507 treatment

TNFα/IFN-γ treatment: Control and MRLC-AA mutant Caco-2 cells were grown on glass coverslips or glass-bottom imaging dishes. Upon reaching 60-70% confluency, cells were treated with 10ng/ml TNFα (R&D Systems, Cat#210-TA) and 10ng/ml Interferon-γ (IFN-γ) (Sigma, Cat#I3275) for 24 hours. ATM1001/ATM3507 treatment: Following 24hrs of TNFα/IFN-γ treatment, cell media was supplemented with tropomyosin inhibitors ATM1001 (2.5μM, 8hrs) or ATM3507 (2.5μM, 8hrs) +/-etoposide (500μM, 8hrs) (Currier et al. 2017; Kee et al. 2018).

### Immunostaining

Caco-2 cells were grown on glass coverslips until they reached 90-100% confluency. The cells were then fixed with 4% paraformaldehyde (PFA, 20min, RT) (Electron Microscopy Sciences, Cat#15710) prepared in cytoskeletal stabilization buffer (10mM PIPES, pH 6.8, 100mM KCl, 300mM sucrose, 2mM EGTA, 2mM MgCl2). This was followed by a 5 minutes permeabilization with 0.25% Triton X-100. For vinculin immunostaining, cells were first pre-permeabilized with an ice-cold permeabilization buffer (0.5% TritonX-100, 10mM PIPES, 50mM NaCl, 3mM MgCl2, 300mM sucrose and 1X protease inhibitor) for 5 min on ice, and, thereafter, fixed with ice cold 50% methanol diluted with acetone (1:1). Fixation was followed by blocking (1h, RT) with 3% bovine serum albumin (BSA/0.01% Tween™20/PBS) (Sigma, Cat#A-7906). Samples were incubated with primary antibodies (overnight, 4°C). Next, the coverslips were washed with 0.01% Tween™20/PBS, followed by incubation with secondary antibody (1h, RT). Coverslips were subsequently washed with 0.01% Tween™20/PBS and mounted with Prolong Gold Antifade reagent with DAPI (Cell Signalling Technologies, Cat#8961).

### Microscopy and image analysis

Images of fixed samples were acquired using Zeiss LSM710 confocal microscope (40x oil objective with 1.3 N.A., 1μm Z-slicing) driven by ZEN software. For quantification of junctional protein intensity, junctional immunofluorescence intensity was measured according to corrected total cell fluorescence (CTCF) analysis in ImageJ. Briefly, a segmented line of 5-10 pixel width was drawn parallelly over the junction, and its integrated density and area was measured. Then, the line was moved on either side of the junction to measure the background mean fluorescence. Next, CTCF was measured through the formula CTCF = integrated density – (area of selected junction x average of background mean fluorescence). For each biological sample, a minimum of 50 junctions were analysed across at least 3 independent repeats and the average junctional intensity was normalised to the control to compare fold-change across the groups.

### Ratiometric vinculin/α-catenin and α-18/α-catenin image processing

To obtain ratiometic images of vinculin/α-catenin and α-18/total α-catenin staining, images were opened in Fiji and max projected (Image −> Stacks −> Z-project… −> Projection type: Max Intensity). α-catenin channel was then processed using Gaussian Blur filter (Process −> Filters −> Gaussian Blur… −> Sigma (Radius): 1) and thresholded to obtain junctional fraction of the staining. Junctional fraction was then selected (Edit −> Selection −> Create Selection −> Make Inverse) and added to ROI. Obtained ROI was then applied to the max projected channels, non-selected space was filled with black colour. Obtained junctional image faction were then divided (Process −> Image Calculator…) creating a 32-bit result ratiometric image, which was then presented as Fire LUT.

### Immunoblotting

Cells were lysed in 1X lysis buffer (50mN Tris-HCl, 10% glycerol, 2% SDS and 0.1% Bromophenol Blue) (98°C, 10 minutes). The cell lysates were resolved in 12% or 14% SDS polyacrylamide gels, followed by transfer to nitrocellulose membrane and block with 5% non-fat milk in 0.01% Tween™20/TBS (1 h, RT). The membranes were incubated with primary antibodies diluted in 3% BSA/0.01% Tween™20/TBS (overnight, 4°C). Thereafter, the membranes were washed with 0.01% Tween™20/TBS and incubated with the respective horse radish peroxidase (HRP)-conjugated secondary antibody diluted in 3% BSA/0.01% Tween™20/TBS (1 h, RT). The blots were visualized with Supersignal West Pico PLUS Chemiluminescent Substrate (Thermo Fisher #34579) and imaged on Biorad Chemidoc.

### Junctional laser ablation

ZO-1-mCherry tagged MRLC-DD mutant and control Caco-2 cells were cultured up to 90-100% confluency in 35-mm glass bottom dishes. Imaging was performed on Zeiss LSM710 confocal microscope (63x oil objective with 1.4 N.A., 3x zoom, 1μm Z-slicing) equipped with 2-photon fully tunable MaiTai eHP DeepSee 760-1040nm laser and a 37°C heating chamber. An oblation ROI, sized 8×30 pixel, was positioned perpendicular to the longer axis of the ZO-1 marked junction. Following acquisition of a first frame, the ROI was ablated using 790nm 2-photon laser at 30-40% transmission power for 10 iterations. Subsequent recoil of the cut junction was recorded for up to 1min with 2sec frame intervals.

To analyse the recoil, the distance *(l)* between the vertices of the ablated junction was tracked in ImageJ with the MTrackerJ plugin and measured as a function of time *(t)*. Each junction was modelled as Kelvin-Voigt fiber and mean values of tracked distances before and after laser ablation (*l(0)* and *l(t)*, respectively) were plotted against time, and calculated by nonlinear regression of the data to the following equation -

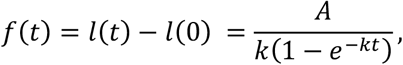

where *l(t)* is defined as the distance between the two vertices at any time point *t*,

*A* is the initial recoil indicating the tensile force at the junction before laser ablation, and *k* is the viscosity coefficient assumed to be constant and calculated as –

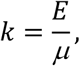

where *E* is the elastic modulus of the junction, and

*μ* is the viscosity coefficient related to viscous drag of the media. *k* values not changing significantly reflect that the initial recoil of the junction is a feature of changes in tensile force alone.

### Etoposide induced apoptosis

Caco-2 cells expressing MRLC-DD or -AA or their respective controls were grown on glass coverslips up to 90-100% confluency and were then treated with 500μM etoposide (AdooQ Bioscience, Cat#A10373) for 8 hours at 37°C. After treatment, cells were fixed with 100% ice-cold methanol for 10 minutes and immunostained for junctional marker ZO-1, apoptotic cell marker cleaved caspase-3 and nuclear stain DAPI. Extrusion efficiency was quantified based on junctional closure of the neighbour cells underneath the apoptotic cell positive for cleaved caspase-3.

### Laser injury induced cell injury

Laser-induced cell injury was performed on Zeiss LSM710 confocal microscope equipped with the 2-photon fully tunable MaiTai eHP DeepSee 760-1040nm laser and a 37°C heating chamber. Double positive ZO-1-mCherry/MRLC-AA or ZO-1-mCherry/control GFP Caco-2 cells were cultured up to 90-100% confluency in 35-mm glass bottom dishes. To induce apoptosis, nuclear micro-irradiation was performed using 10 iterations of 790nm 2-photon laser at 50-70% transmission power (63x oil objective with 1.4 N.A., 1μm Z-slicing). To confirm successful induction of apoptosis, cell media was supplemented with an apoptotic detection marker AnnexinV (Invitrogen, Cat#A23204). Extrusion was considered successful if, (1) injured cell stained positive for AnnexinV and was apically expelled from the monolayer within 60 minutes of laser injury, and (2) the neighbour cells’ junctions closed under it.

### TET-Inducible Puma

Double positive MRLC-DD^PUMA^ or Control^PUMA^ cells were seeded with MRLC-DD and control GFP Caco-2 cells in a 1:100 ratio onto glass coverslips. Once confluent, the co-cultures were treated with 2μg/ml doxycycline (Sigma, Cat#D9891) to initiate apoptosis in the Puma cells. Cells were fixed at 4h, 6h and 8h timepoints and were immunostained for apoptotic marker cleaved caspase-3 and nuclear stain DAPI. Extrusion efficiency was henceforth quantified at each time point. A successful extrusion considered if (1) the Puma-mCherry cell stained positive for cleaved caspase-3, and (2) the apoptotic cell’s nucleus was located above the plane of focus of the healthy neighbour cells in monolayer.

### Statistics

All graphs represented are the mean ± SEM from at least 3 independent repeats. Statistical analyses were performed using GraphPad Prism 9, to obtain p-values with 95% confidence intervals. Unpaired student’s t-test was used to compare datasets between two groups, and either one-way or two-way ANOVA used to compare 3 or more groups, depending on the number of variables.

## Acknowledgements

We thank our colleagues for their support and advice throughout this project. Our work was supported by grants from the National Health and Medical Research Council of Australia (1136592 and 1163462 to A.S.Y; 1079866 and 1100202 to P.W.G and E.C.H); the Australian Research Council (DP220103951 to A.S.Y); Australian Department of Industry, Science and Resources (CRC-P-355 to P.W.G and E.C.H); and by the US Department of Defence (Discovery Grant HT94252310088 to A.S.Y, J.B and J.M.D). Microscopy was performed at the ACRF/IMB Cancer Research Imaging Facility created with the generous support of the Australian Cancer Research Foundation. Additionally, we acknowledge support from the NCCS Bioimaging Centre.

## Supplemental Figure legends

**Figure S1. Characterisation of MRLC-DD Caco-2 line. Related to Figure 1**.

**(A)** Schematic diagram showing the mechanical changes within tissue during apoptotic cell extrusion.

**(B)** Representative images of cells expressing soluble GFP (Ctr) or GFP-tagged phosphomimetic MRLC mutant, MRLC-DD.

**(C)** Representative immunoblot of control GFP or GFP-tagged MRLC-DD expression.

**(D-E)** A representative immunoblot (D) and quantification (E) of total protein levels of non-muscle Myosin IIA in control and MRLC-DD expressing lines.

**(F-G)** A representative immunoblot (F) and quantification (G) of total protein levels of non-muscle Myosin IIB in control and MRLC-DD expressing lines.

**(H-J)** Selected frames (H) from live imaging of 2-photon laser-mediated junction ablation followed by junctional recoil in control and MRLC-DD cells. Cyan line - initial position of junctional vertices, magenta line - actual position of recoiling vertices; (I) Speed of recoil of ablated junctions (data shows one experimental repeat); (J) K-values calculated for ablated junctions in control and MRLC-DD cells.

Scale bars: 15μm. XY panels are maximum projection views of all z-stacks. All data are means ± SEM; ns – not significant calculated from n≥3 independent experiments analysed with unpaired Student’s t-test (D, F, I).

**Figure S2. PUMA mixing experiment. Related to Figure 2**.

**(A)** Schematic diagram showing the experimental design for PUMA mixing experiment from Fig 2F.

**(B)** Representative images of apoptotic PUMA cells upon 8hrs doxycycline (2μg/ml) treatment. Orange line-location of the XZ views.

Scale bars: 15μm. XY panels are maximum projection views of all z-stacks.

**Figure S3. Impact of TNFα on recoil of ablated junctions. Related to Figure 3**.

**(A)** Speed of recoil of ablated junctions in control and TNFα-treated cells (data shows one experimental repeat).

**(B)** K-values calculated for ablated junctions in control and TNFα-treated cells.

Data are means ± SEM; ns – not significant calculated from n≥3 independent experiments analysed with unpaired Student’s t-test.

**Figure S4. TNFα does not enhance apoptotic rates in Caco2 cell line. Related to Figure 4**.

**(A)** Representative images of extrusion efficiency in control and TNFα-treated cell lines. Apoptosis was induced by etoposide treatment (500μM, 8hrs). Orange lines-location of the XZ views; asterisk– retained apoptotic cell; arrowhead – junctional closure underneath the extruded apoptotic cells.

**(B)** Number of apoptotic cells detected in live imaging in control and TNFα-treated cell lines. Apoptosis was induced by etoposide (250μM, 16hrs).

**(C-D)** Quantification (C) and a representative immunoblot (D) of cleaved caspase-3 in control and TNFα-treated cell lines. Apoptosis was induced by etoposide (250μM, 16hrs).

Scale bars: 15μm. XY panels are maximum projection views of all z-stacks. All data are means ± SEM; ns – not significant, **p<0.01, ***p<0.001, calculated from n≥3 independent experiments analysed with one-way ANOVA.

