## Supplementary figures and images for "Pre-existing tissue mechanical hypertension at adherens junctions disrupts apoptotic extrusion in epithelia" (from another version of this preprint: version 1, 2023-09-01)

### Supplementary Figures

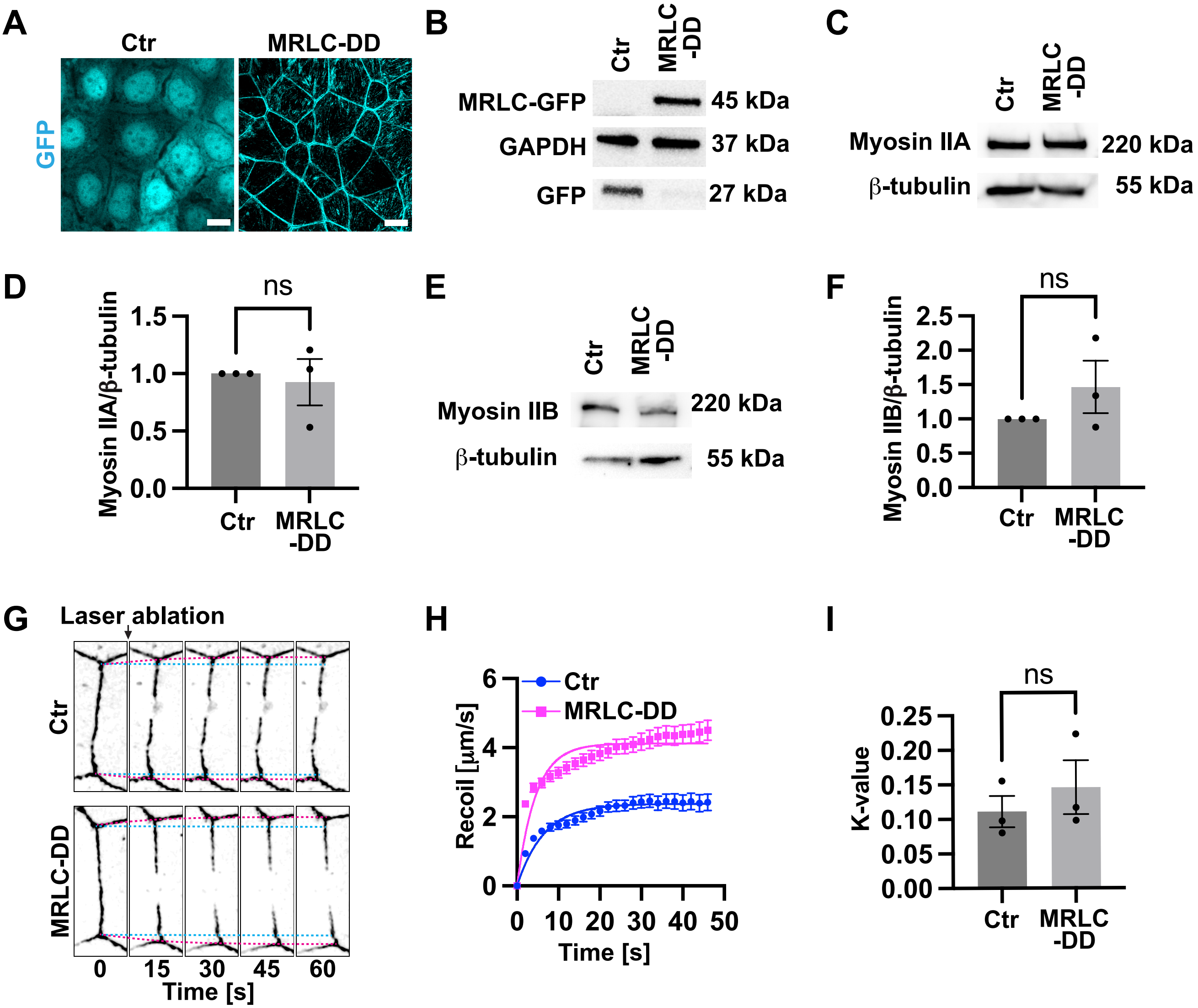

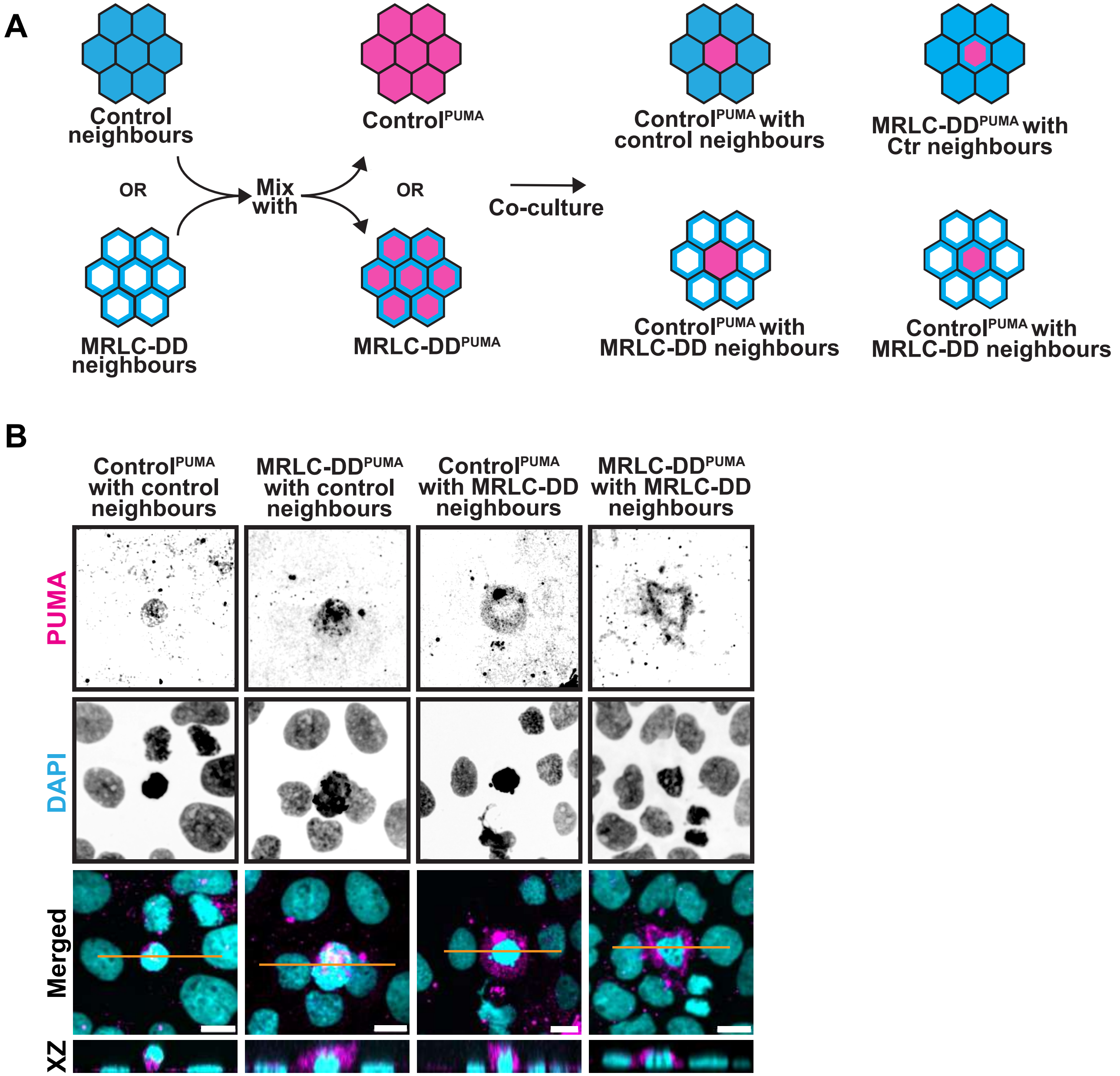

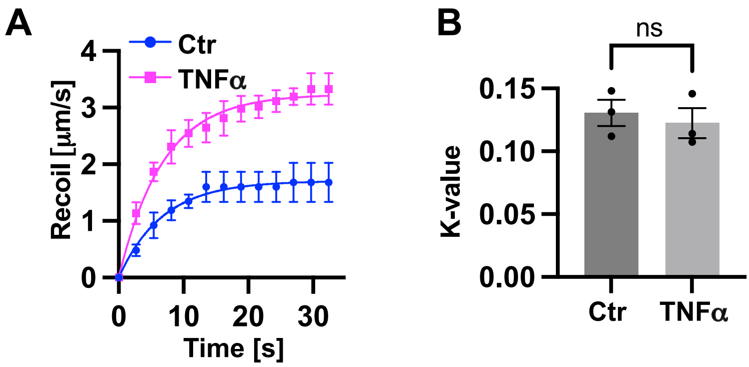

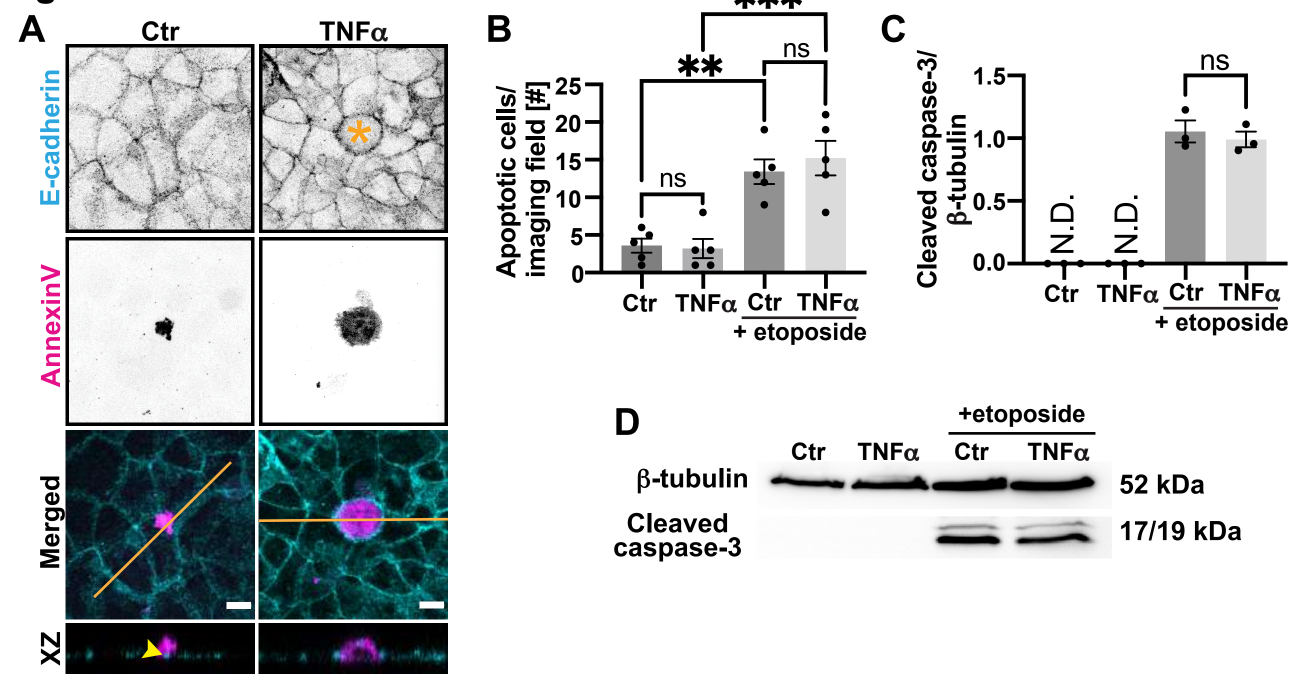
